# Observed Kinetics of Enterovirus Inactivation by Free Chlorine Is Host Cell-Dependent

**DOI:** 10.1101/2022.10.03.509468

**Authors:** Shotaro Torii, Shannon Christa David, Odile Larivé, Federica Cariti, Tamar Kohn

**Affiliations:** Laboratory of Environmental Chemistry, School of Architecture, Civil and Environmental Engineering (ENAC), École Polytechnique Fédérale de Lausanne (EPFL), Lausanne, Switzerland

**Keywords:** Virus, Disinfection, Free chlorine, Host Cells, Inactivation, Water treatment

## Abstract

The virucidal efficacy of disinfectants is typically assessed by infectivity assay utilizing a single type of host cell. Enteroviruses infect multiple host cells via different entry routes, and each entry route may be impaired to a varying extent by a given disinfectant. Yet, it is not known how the choice of host cells for titration affects the observed inactivation kinetics. Here, we evaluated the inactivation kinetics of echovirus 11 (E11) by free chlorine, ultraviolet (UV) irradiation, and heat, using three different host cells (BGMK, RD, and A549). E11 inactivation by free chlorine occurred at a two-fold greater rate when enumerated on BGMK cells compared to RD and A549 cells. Conversely, a comparable inactivation rate was observed for UV and heat independent of the host cell used. Host cell-dependent inactivation kinetics by free chlorine were also observed for echovirus 7, 9 and 13, and coxsackievirus A9, confirming that this phenomenon is not serotype-specific. Inactivation of E11 was partly caused by a loss in host cell attachment, which was most pronounced for BGMK cells, and which may be promoted by a lack of CD55 attachment receptors on this cell type. Additionally, BGMK cells lack a key subunit of the uncoating receptor, β2M, which may further contribute to the differential inactivation kinetic for this cell type. Consequently, inactivation kinetics of enteroviruses should be assessed using host cells with different receptor profiles. This will yield a more complete understanding of the inactivating power of disinfectants targeting the viral attachment and/or uncoating.

**Graphic for Table of Contents (TOC):** 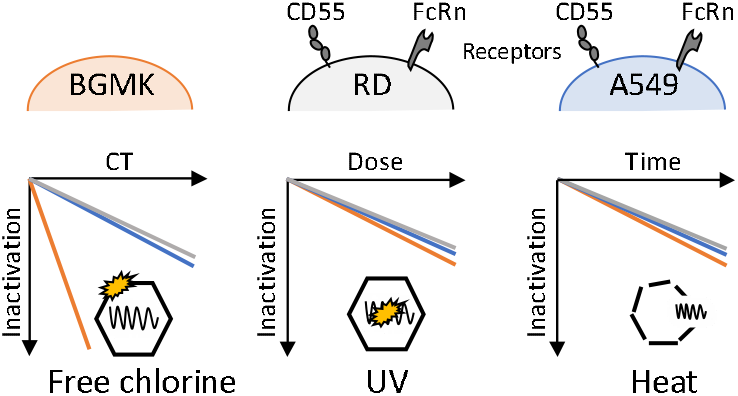

## Introduction

Enteroviruses are non-enveloped positive single-stranded (ss) RNA viruses comprising more than 100 serotypes that infect humans and can cause serious diseases ^1^. These viruses are excreted from the feces of infected persons into the sewage system, and are often detected in wastewater and surface waters ^2^, and have even been reported in tap water after disinfection ^3^. Enteroviruses are therefore included as microbial contaminants in the Draft Contaminant Candidate List 5 published by the US Environmental Protection Agency (USEPA). Due to their ubiquity, enterovirus inactivation during water and wastewater disinfection has been extensively studied ^4–10^.

To evaluate the virucidal efficacy of various disinfectants, methods relying on the infection of a host cell are utilized (e.g., plaque assay and endpoint dilution assay). Buffalo green monkey kidney (BGMK) cells ^11^ are most commonly used for enterovirus titration to assess the efficacy of disinfection in water treatment ^4,7,8,12–17^. This is assumingly because BGMK cells are the most efficient for enterovirus isolation ^18^ and are employed to perform EPA method 1615 ^19,20^ which quantifies “total culturable virus” in water samples. The use of BGMK cells is a *de facto* standard for the measurement of infectious enterovirus in disinfection studies.

However, many enteroviruses are known to infect not only one but multiple types of host cells. For example, one serotype of *Enterovirus*, echovirus type 11 (E11) is also able to infect human rhabdomyosarcoma (RD) ^21^ and human alveolar basal epithelial (A549) cells ^22,23^. The infection starts by virus attachment to host cells. Previous studies have found that the decay-accelerating factor (DAF or CD55) is an attachment receptor for several enteroviruses, including E11 ^24^. While viral attachment to CD55 enhances the efficiency of infection, cells are still permissive to E11 infection even after the suppression of CD55 by gene knockout or by blocking antibodies ^25,26^. Importantly, E11 readily infects BGMK cells which lack CD55 on their surface ^27^, indicating that E11 can attach to host cells via multiple receptors. To trigger entry into the host, the virus must additionally interact with an uncoating receptor, such as neonatal Fc receptor (FcRn) ^26,28^, which is a major histocompatibility complex class I-like protein comprising a heavy α chain and a light β2-microglobulin (β2M) ^29,30^. β2M is also reported to be essential for E11 infection on specific types of host cells (e.g., HEK293T, human osteosarcoma U2OS cell line, and RD cells) ^25,26,28^; however, no studies have investigated the expression of FcRn and its impact on E11 infectivity on BGMK cells.

Disinfectants inactivate viruses by inhibiting one or more viral functionalities, including attachment to cells, entry into cells (i.e., uncoating), and genome replication. The principle mechanism of inactivation differs depending on the disinfectant of interest ^31^; free chlorine and heat mainly impair the attachment function of E11 while ultraviolet irradiation (UV) mainly damages viral RNA, driving the loss of the replicative function ^32,33^. As host cell receptors can be highly variable across cell types, we expect inactivation kinetics to differ depending on the host cells when assessing disinfectants that target the viral attachment or uncoating stage. However, the effect of host cell choice has only been studied for a disinfectant targeting the replicative function (UV irradiation) using adenovirus ^34^.

Here, we assessed the inactivation kinetics of a model enterovirus (E11 Gregory strain) by free chlorine, UV irradiation, and heat, by individual enumeration using three different types of host cells (BGMK, RD, and A549 cells) that are all permissive to enterovirus infection. We also characterized the receptor profiles of these cells to investigate the mechanism of the host cell-dependent inactivation kinetics. Finally, we tested additional serotypes (echovirus 7, 9, 13 (E7, E9, and E13), coxsackievirus A9, B1 (CVA9 and CVB1)) to generalize our findings to other members of the *Enterovirus* genus.

## Materials and Methods

### Cells and Viruses

BGMK and A549 cells were kindly provided by Spiez Laboratory and the Lausanne University Hospital, Switzerland, respectively. RD cells were purchased from the American Type Culture Collection (ATCC) (CCL-136). BGMK cells were grown in Minimum Essential Medium (MEM; Gibco, UK) while A549 and RD cells were grown in Dulbecco’s Modified Eagle Medium (DMEM; Gibco). Both media types were supplemented with 10% fetal bovine serum (FBS; Gibco) and 1% penicillin-streptomycin (P/S; Gibco), and cells were incubated at 37°C with 5% CO_2_.

E11 Gregory strain (ATCC VR-41) and CVB1 environmental isolate (Accession No.: MG845887) ^15^ were propagated on BGMK cells maintained in MEM supplemented with 2% FBS and 1% P/S. E7, E9, E13, and CVA9 were kindly provided by Soile Blomqvist and Carita Savolainen-Kopra (Finnish National Institute for Health and Welfare) and were propagated on RD cells maintained in DMEM supplemented with 2% FBS and 1% P/S. All viruses were allowed to propagate in cells for 3 days, after which the cell flasks were frozen at −80°C and then thawed at room-temperature. The infected cell suspension was centrifuged at 3,000 × g for 15 min. The supernatant was filtered through a 0.45 μm low protein binding durapore membrane (Merck Millipore Ltd., Ireland). 20 mL of the filtrate was then ultracentrifuged at 150,000 × g at 4°C for 3 h (Beckman Coulter, USA) through a 20% (w/v) sucrose cushion. The pellets were resuspended with a 500 μL of phosphate buffer (PB; 1 mM, pH 7.0) (United Chemical Technologies, USA). The suspension was further filtered with a 0.22 μm hydrophilic PTFE membrane (BGB Analytik AG, Switzerland). A 500 μL of PB was further filtered to collect the remained virus on the membrane. The purified stock was stored at 4°C before disinfection experiments. The concentrations of purified stocks were enumerated as detailed below and reported in Table S1 in Supporting Information (SI).

### Enumeration of Infectious Viruses

The concentrations of infectious viruses were quantified by endpoint dilution assay using BGMK, RD, and A549 cells in 96-well plates as previously described ^35^. In brief, the samples were serially diluted 10-fold in the corresponding maintenance medium (MEM or DMEM, both of which were supplemented with 2% FBS and 1% P/S) Then, 150 μL of the diluted samples were inoculated on the host cells, with five replicates per dilution. After incubation at 37□ with 5% CO_2_ for five days, the presence of a cytopathic effect (CPE) in each well was checked by microscopy. The number of positive wells showing CPE for each dilution was counted and converted to adjusted most probable numbers (MPN) using the R package {MPN} ^36^. The lower limit of detection (LoD) was defined as the concentration where CPE was observed at one out of five wells at the lowest dilution, corresponding to 12 MPN/mL in this study.

### Disinfection Experiments

Disinfection experiments were run in duplicate or triplicate for each virus-disinfectant pair tested here. For each run, a total of three time-series samples plus an untreated sample (i.e. sample at time zero), were taken. All experiments were conducted in disinfectant demand-free 1 mM PB (pH 7.0). After the experiment, untreated and disinfected samples were stored at 4□ for a maximum of 24h, then each sample was split into three individual aliquots and was enumerated on each of the three host cell types.

#### Free Chlorine

The free chlorine disinfection experiment was conducted in a batch system. A free chlorine working solution was prepared by diluting sodium hypochlorite (Reactolab SA, Switzerland) in 1 mM PB (pH 7.0). The final free chlorine concentration in the working solution ranged from 0.54 to 0.57 mg L^−1^. The free chlorine concentration was measured by the *N, N*-diethyl-*p*-phenylenediamine (DPD) method ^37^ using a DR300 Chlorine Pocket Colorimeter (Hach Company, USA). Before each run, glass beakers were soaked with >50 mg L^−1^ of sodium hypochlorite overnight to quench the residual chlorine demand. The beakers were rinsed twice with MilliQ water and once with the chlorine working solution. Then, 10 - 50 μL of virus stock solution was spiked into 11.5 mL of the working solution under constant stirring in the quenched beakers. A 500 μL aliquot was collected every 10−50 s (depending on the virus) and mixed with 5 μL of 5,000 mg L^−1^ sodium thiosulfate (Sigma-Aldrich, Germany) to instantly quench the residual free chlorine. The free chlorine concentration in the beaker was measured at the beginning and ten seconds after the collection of the last time-series sample. The decay in free chlorine concentration was less than 12% throughout each run. The chlorine exposure (CT value; concentration of free chlorine multiplied by contact time) for each sample, was given by integration of the time-dependent disinfectant concentration over exposure time, assuming the first-order decay in free chlorine concentration between the two time points.

#### UV

UV irradiation was performed in a collimated beam low-pressure UV system. The UV system comprised an 18 W low-pressure UVC lamp (model TUV T8; Philips, Netherlands), which emitted 253.7 nm light as a quasi-collimated beam ^38^. The fluence rate was determined to be 116-136 μW cm^−2^ by chemical actinometry ^39^. A 5 mL aliquot of 1 mM PB spiked with 30 μL of virus stock was irradiated in a 20 mL beaker with gentle stirring. A 400 μL aliquot was collected every 30 seconds. The collected samples were stored at 4□ until virus enumeration as specified above. The UV exposure (mJ cm^-2^) for each sample was determined as a product of the fluence rate and the corresponding exposure time.

#### Heat Treatment

Heat treatment was conducted in a thermal cycler (GeneAmp PCR system 9700, Applied Biosystems, USA). Five microliters of purified virus stock was spiked into thin-wall PCR tubes containing 45 μL of 1 mM PB. The tubes were incubated at 50□ for either 20s, 40s, or 60s. The incubated tubes were immediately cooled down by placing them on crushed ice, and samples were stored at 4□ as specified above until enumeration.

### Estimation of Inactivation Rate Constants

Inactivation rate constants (*k*_infectivity_) were estimated by fitting a pseudo-first-order model for free chlorine and UV or a first-order model for heat (see Table S2) to the corresponding experimental data, excluding those under the LoD. The rate constants were determined based on the pooled data from all replicates as the slope of ln(*N*/*N*_*0*_) versus disinfectant exposure (free chlorine and UV) or time (heat) by linear least-squares regression, where *N* is the infectious virus concentration at time *T* (MPN mL^−1^), *N*_0_ is the infectious virus concentration at time 0 (MPN mL^−1^), both of which were determined on the corresponding host cells.

### Host attachment of E11 Treated by Free Chlorine

The capacity of E11 to bind to host cells was quantified, with a slight modification of our previous report ^33^. Untreated and disinfected virus samples were diluted 10-fold in maintenance medium. 500 μL of diluted virus samples were added to cell monolayers (BGMK, RD, or A549) in 12-well plates for 1 h at 4□. The cell monolayers were then washed three times with 300 μL of phosphate-buffered saline (PBS, pH 7.4) per well to remove unbound viruses and free RNA. 140 μL of PB was added to each well and samples were stored at −20□ for a maximum of 24 h prior to RNA extraction. After thawing, 560 μL of AVL buffer with carrier RNA, both taken from the QIAamp Viral RNA Mini Kit (QIAGEN, Germany), were added to each well and incubated for 10 min at room temperature, to lyse the bound viruses. The lysate was processed according to the manufacturer’s instruction to obtain RNA extracts in 40 μL of ultrapure water. The RNA concentration was measured by reverse transcription digital PCR (RT-dPCR) using reported primers and probe ^40,41^ on QIAcuity dPCR 2-plex platform (QIAGEN) as detailed in the SI.

The observed rate constants of attachment loss (*k*_obs_attachment_) were determined based on the pooled data from triplicate experiments as the slope of ln(*GC*/*GC*_0_) versus CT value by linear least-squares regression, where *GC*_0_ and *GC* are the number of bound viruses (measured as genome copies) before and after chlorine exposure, respectively. Note that the difference in observed genome copy numbers between untreated and disinfected samples stems from a reduction in bound viruses, as well as from the decay of the targeted PCR segment due to exposure to free chlorine ^33^. To determine the net rate constants for attachment loss (*k*_attachment_), *k*_obs_attachment_ was thus corrected by the observed decay of the PCR-target (*k*_PCR-target_), which was measured in sample aliquots not included in the binding assay (see Eq.(1)).

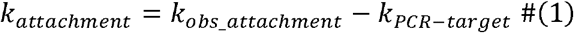

### Flow Cytometric Analysis of Cell Receptors

Cells were washed with PBS, detached by 0.05% trypsin-EDTA (Gibco), and pelleted in a 96-well U-bottom plate (Thermofisher Scientific, USA) at 10^5^ cells per well by centrifugation at 400 × g for 2 min. Cells were washed twice in PBS and then stained with 1 μg/mL of DAPI (Sigma-Aldrich) for 15 min at room temperature in the dark. After being washed twice by staining buffer (PBS with 1% of bovine serum albumin), cells were incubated with Fc receptor blocking solution (BD Pharmingen, USA) for 15 min at room temperature in the dark. Subsequently, cells were stained with FITC-conjugated anti-human CD55 antibody (Biolegend, USA) (2 μg/mL) and APC-conjugated anti-human β2-microglobulin (β2M) antibody (Biolegend) (0.25 μg/mL), or isotype controls (FITC-conjugated mouse IgG1 (Biolegend) and APC mouse IgG1-conjugated (Biolegend) for 20 min at 4□ in the dark. Stained cells were washed twice with PBS and resuspended with 200 μL of PBS prior to immediate acquisition. Data acquisition for all samples was performed on a Gallios flow cytometer (Beckman Coulter) at the EPFL Flow Cytometry Core Facility, with a minimum of 10,000 cells acquired per sample. Acquired data were analyzed in FloJo software version 10.8.0. Cell doublets were excluded by single cell gating, and single cells were then gated based on viability (DAPI^-^) prior to analysis of cell receptor expression (CD55^+^ / β2M^+^). Unstained cells and those stained with isotype control antibodies were used to set negative gates for fluorescence for each cell type.

### Statistical Analyses

All statistical analyses were implemented in R-4.1.2 ^42^. Linear least-squares regression was performed with the {lm} function to estimate inactivation rate constants from the disinfection experiments. Inactivation rate constants of the different host cells were compared by analysis of covariance (Type III) using {Anova} function in the car package ^43^. If a significant effect of host cells was observed, a post-hoc analysis was performed using {emtrends} function in the emmeans package ^44^ to determine which pairs of host cells are significantly different. The alpha-type error was set at 0.05.

## Results and Discussion

### Observed Kinetics of E11 Inactivation Enumerated by Different Host Cells

Prior to conducting inactivation experiments, we ensured that all three host cells were permissive to E11 infection. We found that each cell type produced high E11 titers, (ranging from 6.9 – 7.9 log_10_ MPN/mL; Table S1), suggesting that all the cells were readily infected by E11.

To assess the impact of host cells on the observed kinetics of E11 inactivation, untreated and disinfected samples were enumerated individually on the three different host cells included in this work. The resulting inactivation curves are shown in Figure 1. The observed inactivation efficiencies for free chlorine were higher for E11 samples enumerated with BGMK cells compared with RD and A549 cells. For example, at a CT 0.18 mg min L^-1^, E11 Gregory was inactivated by 2.8 log_10_ on BGMK cells, compared with the lower 1.2 and 1.3 log_10_ seen on RD and A549 cells, respectively. (Figure 1A). The observed inactivation rate constants (*k*_infectivity_) were 25.0, 12.6, and 14.6 mg^-1^ min^-1^ L for BGMK, RD, A549 cells, respectively (Table 1). The inactivation rate constants for BGMK cells were 2.0- and 1.7-fold higher than those for RD and A549 cells, respectively, and these differences were statistically significant (p-value: 9.5 × 10^−5^ (BGMK - RD) and 1.0 × 10^−3^ (BGMK - A549)). On the other hand, the choice of host cell only led to small and not statistically significant differences in inactivation kinetics of E11 treated by UV or heat (Figure 1B, C). These results demonstrate that the observed inactivation kinetics of E11 depend on the host cells used for enumeration, though the magnitude of the host cell effect depends on the disinfectant.

**Table 1.** Rate constants for the inactivation (*k*_infectivity_) and loss of attachment (*k*_attachment_) after treatment of E11 by free chlorine

| Host cells | $k_{\text{infectivity}}$ (mg<sup>-1</sup> min<sup>-1</sup> L) | $k_{\text{attachment}}$ (mg<sup>-1</sup> min<sup>-1</sup> L) |
| --- | --- | --- |
| BGMK | 25.0 | 14.1 |
| RD | 12.6 | 9.0 |
| A549 | 14.6 | 8.9 |

**Figure 1.**
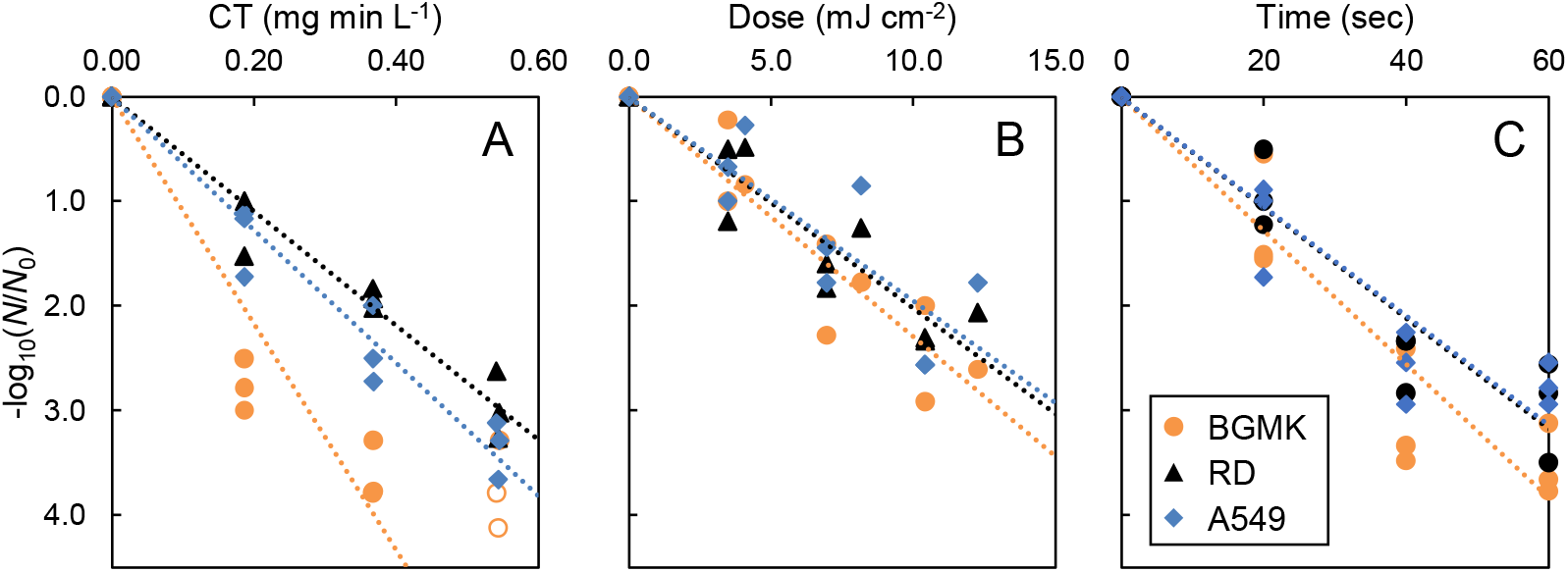
Inactivation of E11 by free chlorine (A), UV (B), and heat (C) measured on BGMK (orange circles), RD (black triangles), and A549 (blue diamonds) cells. Dashed lines show each regression line. Empty symbols indicate right-censored data and are plotted at the value of −log_10_(*LoD*/*N*_0_).

### Potential Mechanism of Host Cell-Dependent Kinetics of Inactivation by Free Chlorine

Free chlorine was previously shown to inhibit the ability of E11 to attach to BGMK cells ^33^. We therefore investigated if the host cell-dependence of E11 inactivation kinetics could be explained by a differential effect of free chlorine on host attachment to the three cell types studied. Figure S1 shows the observed loss of attachment as a function of CT values, and Table 1 shows the rate constants for loss of attachment (*k*_attachment_). The *k*_attachment_ was 14.1, 9.0, and 8.9 mg^-1^ min^-1^ L for BGMK, RD, and A549 cells. This demonstrates that binding capability to BGMK cells was affected most strongly.

This finding can be rationalized by considering the receptor profiles of the three host cells. A flow cytometry analysis was used to characterize the presence of CD55, an attachment receptor, and β2M, a key subunit of the uncoating receptor FcRn, on each of the host cells. Expression of these receptors on BGMK, RD, and A549 cells is shown in Figure S2 and summarized in Table 2. The attachment receptor CD55 was expressed on RD and A549 cells but was not detected on BGMK cells. This confirms a previous finding, which reported that BGMK cells were negative for CD55 while RD cells were positive ^27^. Similarly, β2M was detected on RD and A549 cells, but not on BGMK cells. Expression of β2M on RD cells is consistent with a previous finding ^30^. These results suggest that both RD and A549 cells can interact with E11 via CD55 and β2M while BGMK cells cannot. Given that BGMK cells replicated the untreated E11 as efficiently as the other two host cells (Table S1), BGMK cells must possess alternative attachment and uncoating receptors for E11. This further implies that E11 uses different motifs on the viral capsid to interact with host cell receptors on BGMK cells than on RD or A549 cells. A preference of free chlorine for certain proteins of viral capsids has previously been reported for phages ^31,45,46^. The greater effect of free chlorine on host attachment to BGMK cells may thus stem from preferential oxidation of the binding motif for BGMK compared to CD55.

**Table 2.** Expression of receptors on each of the host cells

| Host cells | CD55 | β2M |
| --- | --- | --- |
| BGMK | Negative | Negative |
| RD | Positive | Positive |
| A549 | Positive | Positive |

The difference in *k*_attachment_ between BGMK and the other two host cells alone, however, cannot explain the much larger difference in the *k*_infectivity_ (Table 1). We hypothesize that the rate of uncoating is also more affected in viruses enumerated on BGMK cells compared to the other two cell types. Given the absence of β2M on BGMK cells (Table 2), E11 must use a different receptor and hence a different host interaction site to successfully uncoat in BGMK cells. If this host interaction site is more susceptible to degradation by chlorine, then this would lead to a faster loss of the uncoating function between viruses enumerated on different host cells. This hypothesis could not be tested in this study, because we were not able to quantitatively measure virus internalization. A published assay ^26^ was tested but was not able to separate between truly internalized and merely bound viruses (data not shown) in our experimental conditions.

We do not expect the replicative function to be differentially affected in different host cells. Loss of the replicative function can be regarded as the loss of intact RNA that can be replicated and transcribed by cellular protein synthesis machinery. Given that the repair of ssRNA within the host cells is unlikely, the intactness of RNA is dependent on virus and treatment and independent of host cells. In fact, for UV, which dominantly damages E11 replicative function ^33,47^, a similar *k*_inactivation_ was observed regardless of host cells, suggesting that a loss of replicative function affects all host cells equally.

Similar to free chlorine, heat was reported to strongly affect *k*_attachment_ of E11 ^33^. Interestingly, the choice of host cell did not influence virus inactivation kinetics by heat. A potential hypothesis is a difference in the damaging mechanism between free chlorine and heat treatment. Previous studies found that mild heat treatment either causes the capsid to dissociate into pentametric subassemblies ^48^ or induces conformational rearrangement ^49^, both of which release enteroviral RNA to the outside of the capsid ^48,49^. Disassembled or rearranged capsids would not be expected to interact with any host cell receptors, such that the attachment of E11 to all host cells, and hence the infectivity of E11, would decline at an equal rate.

### Effect of Host Cells on the Inactivation Kinetics of Other Serotypes of the *Enterovirus* Genus

A panel of enteroviruses (CVA9, CVB1, E7, E9, and E13), which were reported to be prevalent in European sewage ^50^, was tested for their ability to infect the three host cells (Table S1). The results suggested that these viruses can propagate in BGMK cells as well as RD and A549 cells, despite the lack of CD55 and β2M. Subsequently, the five viruses were examined for host cell-dependence in their inactivation kinetics by free chlorine. Inactivation curves were determined for each combination of the virus and the host cells (Figure 2). The effect of host cells on observed inactivation rate constants was significant for all the tested viruses, except for CVB1. The effect size ranged from a 1.3-to a 3.5-fold change in *k*_infectivity_, depending on the serotype. The observed inactivation on BGMK cells was significantly faster than that on RD and A549 cells for CVA9 (p-value: 4.8 × 10^−5^ (BGMK-RD) and 3.8 × 10^−2^ (BGMK-A549)), E9 (p-value: 3.4 × 10^−4^ (BGMK-RD) and 3.8 × 10^−2^ (BGMK-A549)), and E13 (p-value: 2.2 × 10^−4^ (BGMK-RD) and 1.1 × 10^−3^ (BGMK-A549)). A significant difference between RD and A549 cells was only observed for E7 (p-value: 6.9 × 10^−3^).

**Figure 2.**
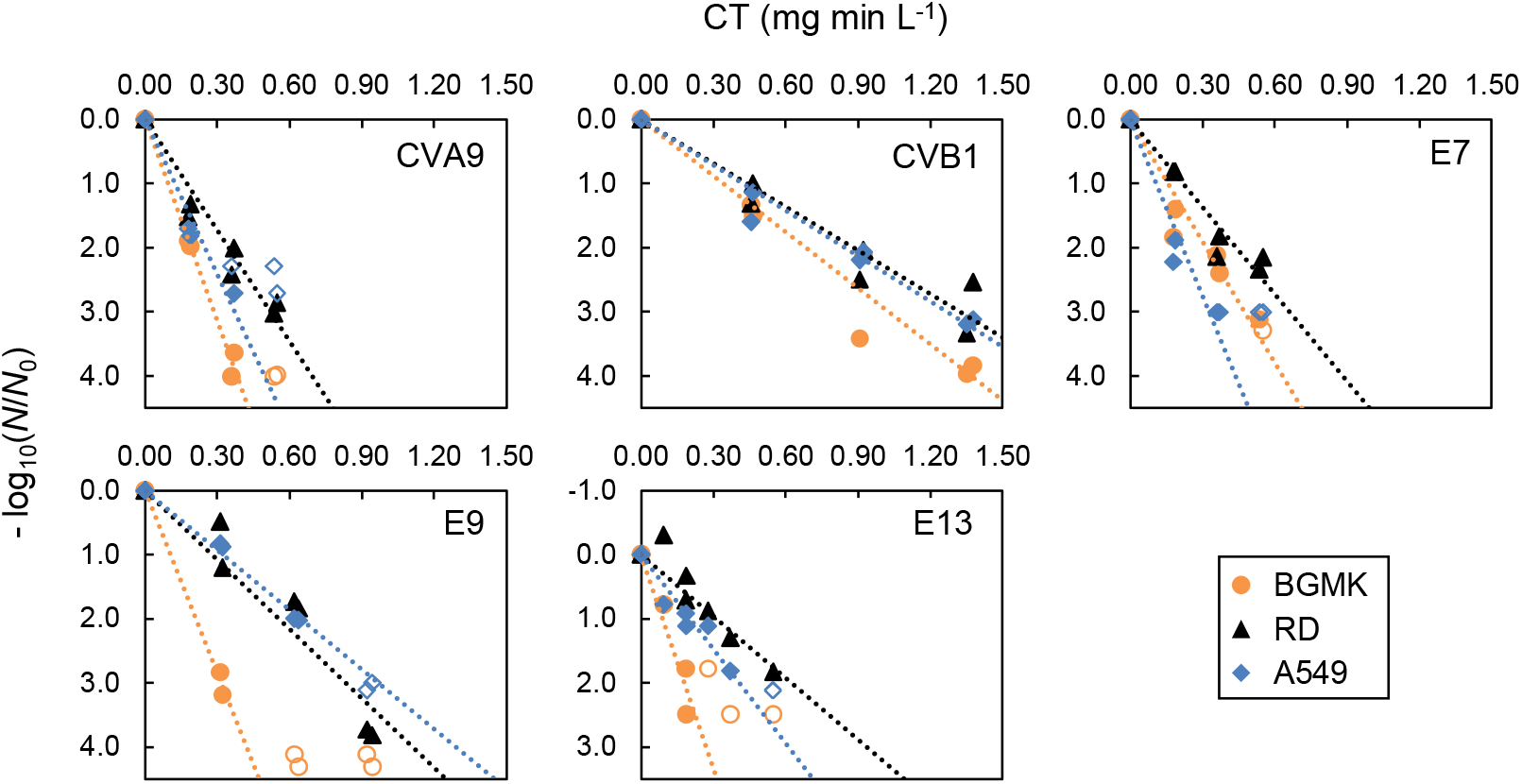
Inactivation of a panel of enteroviruses by free chlorine determined by enumeration on BGMK (orange circles), RD (black triangles), and A549 (blue diamonds) cells. Dashed lines show each regression line. Empty symbols indicate right-censored data and are plotted at the value of −log_10_(*LoD*/*N*_0_).

We attempted to rationalize these observations by considering the receptor profiles of the three host cells (Table S3) and the receptor usage of each serotype (Table S4). CVA9 and E9 were reported to use αVβ3 as an attachment receptor, which is expressed on all cell types used. This attachment receptor can thus not be implicated as the cause for host cell-dependent inactivation kinetics. These viruses furthermore use FcRn for uncoating, which is absent from BGMK cells. As discussed above for E11, CVA9 and E9 must thus rely on an alternative uncoating receptor on BGMK cells, which may lead to the observed increase in sensitivity to free chlorine. E13 was reported to use CD55 as an attachment receptor and FcRn as an uncoating receptor, and thus has the same receptor usage as E11. It is then not surprising that this virus exhibited a similar host cell-dependence in its inactivation kinetics as E11. Interestingly, however, the same receptors are also used by E7, which did not exhibit significantly faster inactivation kinetics when using BGMK cells as the host. This may be rationalized by reports that E7 can also enter cells by clathrin-mediated endocytosis ^51^. The significant difference between inactivation rates measured on RD and A549 cells for E7, however, remains to be investigated. Finally, further investigation is also required to understand the comparable inactivation rates of CVB1 on all host cells. A possible explanation is this virus’ use of the coxsackievirus and adenovirus receptor (CAR), which works bi-functionally as an attachment and uncoating receptor for Group B coxsackieviruses^52^. CAR was reported to be expressed on all three host cells used herein, though the expression level is reported differently among studies ^27,53^. The similarity of the major infection route may explain the similar rate constants among the host cells tested here.

In summary, the host cell-dependent inactivation kinetics for free chlorine was observed for several serotypes of the *Enterovirus* genus. Further research is needed to conclusively determine if the magnitude of the difference is associated with receptor usage of the tested viruses and the availability of attachment and uncoating receptors expressed by each of the host cells.

### Environmental Implications

This work revealed host cell-dependent kinetics of enterovirus after inactivation by free chlorine and suggested potential mechanistic contributions of host cell receptor profiles to this phenomenon. Furthermore, data indicate that host cell-dependent inactivation kinetics are observed for a range of serotypes of the *Enterovirus* genus.

To date, the classic method for assessing the virucidal efficacy of disinfectants is to select any single cell type permissible to infection by the tested virus. Our results caution against this approach, and instead encourage the use of multiple cell types with different receptor profiles. This would reduce the risk of inactivation efficiencies being over- or underestimated, especially for disinfectants damaging the capsid in a selective manner (e.g., chlorine dioxide ^31^). Moreover, an appropriate rationale is necessary to select the host cells used to evaluate disinfection efficiency. For example, this study shows that RD cells can provide more conservative estimates of inactivation efficiency (Figure 2) and observed infectious concentration for echoviruses than BGMK cells (Table S1). A possible approach more representative of human infection may be the use of host cells closer to the gastrointestinal tract (e.g., Caco-2 and HT29 cells), or enteroids ^54^ as an *ex vivo* model of the human intestinal epithelium. Also, *in vivo* disinfection experiments (e.g. those performed by Zhang et al. ^55^) compared with *in vitro* experiments could aid in understanding the meaning of observed inactivation efficiency. Collectively, these new approaches will improve our interpretation of the observed inactivation *in vitro* and understand “true” inactivation kinetics by disinfection.

## Acknowledgment

This work was funded in part by the JSPS Overseas Challenge Program for Young Researchers, the Young Researchers Exchange Programme between Japan and Switzerland (EGJP_04-042020), a JSPS Overseas Research Fellowships to S.T., and by EPFL discretionary funds. We thank the EPFL Flow Cytometry Core Facility for access to instruments and technical help. We also thank Soile Blomqvist and Carita Savolainen-Kopra (Finnish National Institute for Health and Welfare) for providing environmental enterovirus isolates.

## Supporting Information

The Supporting Information is available at XXXXXX.

- RT-dPCR methods used; Infectious virus concentration of virus stocks enumerated on each of host cells; Inactivation models; Expression of receptors for each of host cells; reported receptors for enteroviruses; CD55 and β2M expression on BGMK, RD, and A549 cells; Decay of virus functions with free chlorine.

## Data availability

All data discussed in this manuscript will be made available on zenodo upon acceptance of the manuscript.

