## Supplemental Information for "Observed Kinetics of Enterovirus Inactivation by Free Chlorine Is Host Cell-Dependent"

Contents (9 pages, 1 Text section, 4 Tables, and 2 Figures)

**Text section**

S1. RT-dPCR analysis

**Tables**

Table S1 Infectious virus concentration of virus stocks enumerated on each type of host cell.

Table S2 Inactivation models

Table S3 Expression of receptors for each of the host cells.

Table S4 Reported attachment and uncoating receptors for enteroviruses

**Figures**

Figure S1 Decay of virus functions with free chlorine.

Figure S2 CD55 and β2M expression on BGMK, RD, and A549 cells, determined by flow cytometry.

### S1. RT-dPCR analysis

A RT-dPCR assay for enterovirus RNA quantification was optimized adapting the thermal cycling conditions and primers and probe concentrations from a previously described RT-qPCR assay^1,2^. The assay was performed in 40 μL reaction mixtures using the QIAcuity One-Step Viral RT-PCR Kit (QIAGEN) and 26k 24-well Nanoplates (QIAGEN). The 26k QIAcuity 24-well Nanoplates are microfluidic dPCR plates including 24wells/plate and 26000 partitions/well. PCRs occur in each partition at a volume of 0.91 nL. The enterovirus RT-dPCR mixture contained 10 μL of 4X One-Step Viral RT-PCR Master Mix, 1000 nM forward primer (5’- CCTCCGGCCCCTGAATG -3’), 1000 nM reverse primer (5’- ACCGGATGGCCAATCCAA -3’), 500 nM probe (5’- FAM-CGGAACCGACTACTTTGGGTGTCCGT-TAMRA -3’), 0.4 μL of 100× Multiplex Reverse Transcription Mix, 5.6 μL of DNase and RNase free water, and 15 μL of template RNA. As variation of technical replicates during optimization phase appeared negligible, a singlicate of RT-dPCR was used for each sample. The nanoplate was then loaded onto the QIAcuity One, 2-plex Device (Qiagen). The thermal cycles include an RT step at 55 °C for 60 min, 95 °C for 5 min for enzyme activation, and followed by 45 cycles of denaturation (95 °C for 15 s) and annealing/extension (at 58 °C for 60 s). In each run, a negative control (no template) and a positive control (i.e., a gBlock containing the target sequence (Integrated DNA Technologies, Coralville, IA, USA)) were included. The final imaging step was made by reading in the FAM channel. Data were analyzed using QIAcuity Software Suite 2.0.20 (Qiagen). Quantities were expressed as genome copy (gc) per microliter of the reaction mixture. The RT-dPCR assays were performed using automatic settings for the threshold and baseline. The limit of quantification (LoQ) was determined by measuring six replicates of 12.5, 25, 50, and 100 gc/reaction using the gBlock. The LoQ was defined as the lowest value where the relative standard deviation was smaller than 35%, given that allowable variation, is proposed as a relative standard deviation from 25 to 35%. The LoQ of the assay was determined to be 12.5 gc/reaction, corresponding to 0.315 gc/μL in the reaction mixture.

Table S1 Infectious virus concentration of virus stocks enumerated on each type of host cell. The measurement was performed in triplicates for E11 and in duplicates for the others. Data presented is the mean from replicate measurements.

| Serotype | BGMK  (log_10_ MPN mL^-1^) | RD  (log_10_ MPN mL^-1^) | A549  (log_10_ MPN mL^-1^) |
| --- | --- | --- | --- |
| E11 | 7.2 | 7.9 | 6.9 |
| CVA9 | 8.1 | 8.1 | 6.6 |
| CVB1 | 8.5 | 8.7 | 8.2 |
| E7 | 7.0 | 7.7 | 6.8 |
| E9 | 7.7 | 8.0 | 6.5 |
| E13 | 5.6 | 6.5 | 5.5 |

Table S2 Inactivation models

| Disinfectant | Inactivation model |
| --- | --- |
| Free chlorine | $\frac{N}{N_{0}}=e^{-k_{inactivation}CT} \left( 1 \right)$ |
| UV | $\frac{N}{N_{0}}=e^{-k_{inactivation}ET} \left( 2 \right)$ |
| Heat | $\frac{N}{N_{0}}=e^{-k_{inactivation}T} \left( 3 \right)$ |

*N* is the infectious virus concentration at time *T* (MPN mL^−1^), *N*_0_ is the infectious virus concentration at time 0 (MPN mL^−1^), *k*_inactivation_ is the inactivation rate constant (mg^−1^ min^−1^ L or mJ^−1^ cm^2^ or sec^-1^), *C* is the free chlorine concentration (mg L^-1^), *E* is the fluence rate of UV (mW cm^-2^), *T* is the exposure time (sec or min).

Table S3 Expression of receptors for each of the host cells, based on published literature (αVβ3, αVβ6, CAR) and our experimental data (CD55, β2M). The number of + signs indicates the relative degree of receptor expression. A “-“ sign indicates no expression. The sign “+/-“ indicates that the expression was inconsistently reported in the literature. The expression of CD55 and β2M, a key component of FcRn, were determined in this study.

|  | CD55 | β2M | αVβ3 | αVβ6 | CAR |
| --- | --- | --- | --- | --- | --- |
| BGMK | - | - | ++ ^3^ | +/- ^3,4^ | ++ ^5^ |
| RD | ++ | ++ | + ^3^ | - ^3^ | +/- ^5,6^ |
| A549 | ++ | ++ | + ^3^ | + ^3^ | ++ ^7^ |

Table S4 Reported attachment and uncoating receptors for enteroviruses

| Serotype | Attachment receptor | Uncoating receptor |
| --- | --- | --- |
| E11 | CD55 ^8^ | FcRn ^9^ |
| CVA9 | αVβ3 ^10^, αVβ6 ^4^ | FcRn ^9^ |
| CVB1 | CD55 ^11^, CAR ^12,13^, αVβ6 ^14^ | CAR ^12,13^ |
| E7 | CD55 ^8^ | FcRn ^9^ |
| E9 | αVβ3 ^15^ | FcRn ^9^ |
| E13 | CD55 ^8^ | FcRn ^9^ |

| A  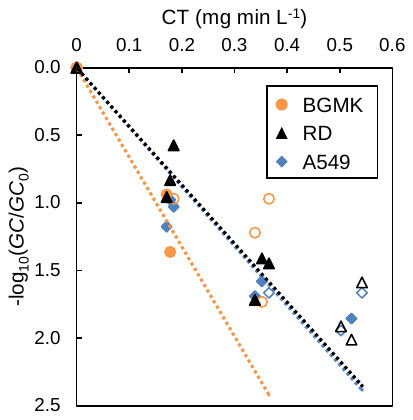 | B  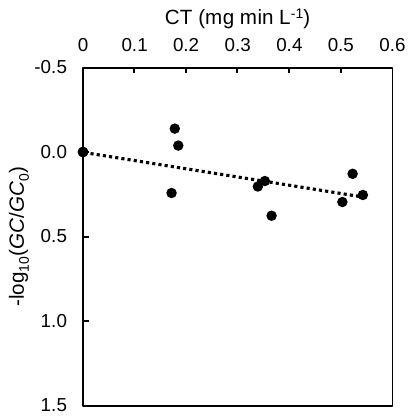 |
| --- | --- |

Figure S1 Decay of virus functions with free chlorine. (A) The observed loss of attachment for each host cells as a function of CT values. The data of BGMK, RD, and A549 cells are shown by orange circles, black triangles, and blue diamonds, respectively. Empty symbols indicate right-censored data and are plotted at the value of -log_10_(*LoQ*/*GC*_0_). The rate constant for the observed loss of attachment (*k*_obs_attachment_) was estimated from these regressions. (B) Observed degradation of the PCR-target as a function of CT values. The rate constant for the integrity loss of the PCR-target (*k*_PCR-target_) was estimated from the regression.

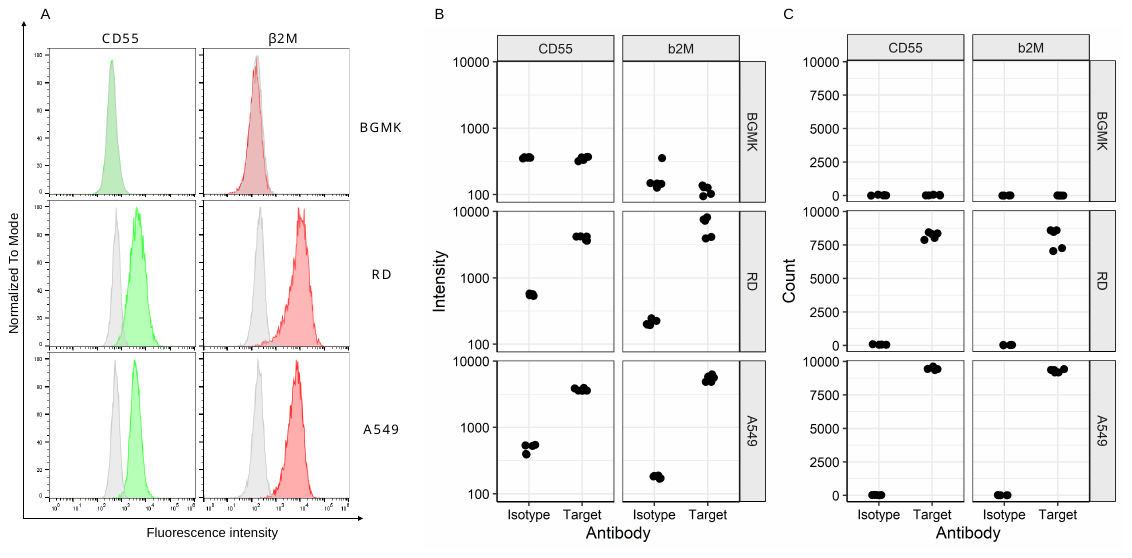

Figure S2 CD55 and β2M expression on BGMK, RD, and A549 cells, determined by flow cytometry. (A) Fluorescent intensity histograms of representative samples for each cell type. For the detection of CD55 and β2M, monoclonal anti-CD55 (FITC) and monoclonal anti-β2M (APC) antibodies were used (green histogram and red histogram, respectively). Fluorophore conjugated IgG1 antibodies were used as isotype controls (grey histogram) in each case. (B). The geometric mean fluorescence intensity for FITC (CD55) and APC (β2M) of single cells that were pre-gated on viability (DAPI -). (C) Total cell counts for CD55^+^ and β2M^+^ cells, compared to cells stained with isotype control antibodies.
